# Variants in the degron of *AFF3* cause a multi-system disorder with mesomelic dysplasia, horseshoe kidney and developmental and epileptic encephalopathy

**DOI:** 10.1101/693937

**Authors:** Norine Voisin, Rhonda E. Schnur, Sofia Douzgou, Susan M. Hiatt, Cecilie F. Rustad, Natasha J. Brown, Dawn L. Earl, Boris Keren, Olga Levchenko, Sinje Geuer, David Amor, Alfredo Brusco, E. Martina Bebin, Gerarda Cappuccio, Joel Charrow, Nicolas Chatron, Gregory M. Cooper, Elena Dadali, Julien Delafontaine, Ennio Del Giudice, Ganka Douglas, Tara Funari, Giuliana Giannuzzi, Nicolas Guex, Delphine Heron, Øystein L. Holla, Anna C.E. Hurst, Jane Juusola, David Kronn, Alexander Lavrov, Crystle Lee, Else Merckoll, Anna Mikhaleva, Jennifer Norman, Sylvain Pradervand, Victoria Sanders, Fabio Sirchia, Toshiki Takenouchi, Akemi J. Tanaka, Heidi Taska-Tench, Elin Tønne, Kristian Tveten, Giuseppina Vitiello, Tomoko Uehara, Caroline Nava, Binnaz Yalcin, Kenjiro Kosaki, Dian Donnai, Stefan Mundlos, Nicola Brunetti-Pierri, Wendy K. Chung, Alexandre Reymond

## Abstract

The ALF transcription factor paralogs, *AFF1, AFF2, AFF3* and *AFF4*, are components of the transcriptional super elongation complex that regulates expression of genes involved in neurogenesis and development. We describe a new autosomal dominant disorder associated with *de novo* missense variants in the degron of AFF3, a nine amino acid sequence important for its degradation. Consistent with a causative role of *AFF3* variants, the mutated AFF3 proteins show reduced clearance. Ten affected individuals were identified, and present with a recognizable pattern of anomalies, which we named KINSSHIP syndrome (KI for horseshoe <u>KI</u>dney, NS for <u>N</u>ievergelt/<u>S</u>avarirayan type of mesomelic dysplasia, S for <u>S</u>eizures, H for <u>H</u>ypertrichosis, I for <u>I</u>ntellectual disability and P for <u>P</u>ulmonary involvement), partially overlapping the *AFF4* associated CHOPS syndrome. An eleventh individual with a microdeletion encompassing only the transactivation domain and degron motif of *AFF3* exhibited overlapping clinical features. A zebrafish overexpression model that shows body axis anomalies provides further support for the pathological effect of increased amount of AFF3 protein.

Whereas homozygous *Aff3* knockout mice display skeletal anomalies, kidney defects, brain malformation and neurological anomalies, knock-in animals modeling the microdeletion and the missense variants identified in affected individuals presented with lower mesomelic limb deformities and early lethality, respectively.

Transcriptome analyses as well as the partial phenotypic overlap of syndromes associated with *AFF3* and *AFF4* variants suggest that ALF transcription factors are not redundant in contrast to what was previously suggested

## Introduction

The *AFF1* (AF4/FMR2 family member 1, a.k.a AF4), *AFF2* (a.k.a *FMR2*), *AFF3* (a.k.a *LAF4*) and *AFF4* genes encode members of the ALF (<u>A</u>F4/<u>L</u>AF4/<u>F</u>MR2) family. These transcription factors share five highly conserved domains starting from the amino terminus: (i) an N-terminal homology domain (NHD); (ii) the hallmark ALF domain, which interacts with Seven In Absentia Homolog (SIAH) ubiquitin ligases through the [xPxAxVxPx] degron motif^1, 2^ and thus regulates protein degradation mediated by the proteasome pathway; (iii) a serine-rich transactivation domain^3^; (iv) a bipartite nuclear localization sequence (NLS); and (v) a C-terminal homology domain (CHD)^4, 5^. *AFF1*, *AFF3*, and *AFF4* have each been identified as fusion partners of the mixed-lineage leukemia (*MLL*) gene involved in acute pediatric leukemias^3^. They are part of the super elongation complex^6^ implicated in transcription of a set of genes, among them histones, retinoid signaling and *HOX* genes involved in neurogenesis and several other developmental processes (e.g. *Hoxa1, Cdx11* and Cyp26a1^6, 7^). Mutations of the fruit fly ALF orthologous gene *lilliputian* (*lilli*) were shown to prevent neuronal differentiation and to decrease cell growth and size^8, 9^. Silencing of *AFF2* by CGG repeat expansion is associated with FRAXE intellectual disability syndrome^10^ (OMIM #309548), whereas hypermethylation of a mosaic CGG repeat expansion in the promoter of *AFF3*, which leads to its silencing in the central nervous system, was associated with a cytogenetic fragile site (FRA2A) and intellectual disability in three families^11^. AFF3 is also known for regulating the expression of imprinted genes^12, 13^ such as *XIST* through binding to differentially methylated regions^14^. An individual carrying a 500kb microdeletion within the *AFF3* locus and presenting with skeletal dysplasia and encephalopathy was described^15^.

Six *de novo* missense variants in *AFF4* were recently linked with CHOPS (<u>C</u>ognitive impairment and coarse facies, <u>H</u>eart defects, <u>O</u>besity, <u>P</u>ulmonary problems, <u>S</u>hort stature and skeletal dysplasia) syndrome^16, 17^ (OMIM#616368). They were suggested to act through reduced clearance of AFF4 by SIAH, a hypothesis supported by the fact that surviving adult *Aff4* null mice have only azoospermia and no features of CHOPS syndrome. However, a majority of *Aff4*^−/−^ embryos died *in utero* with severely shrunken alveoli of the lung^18^. Upregulation of *AFF4* resulted in dysregulation of genes involved in skeletal development and anterior/posterior pattern formation such as *MYC, JUN, TMEM100, ZNF711* and FAM13C^16^. These molecular changes were proposed to impair complex function and lead to cohesinopathies associated with the clinical phenotypes seen in the eleven reported individuals with CHOPS and in Cornelia de Lange syndrome (CdLS; OMIM #122470)^16, 17^.

Here we describe 10 individuals with *de novo* missense variants in the *AFF3* gene and a recognizable pattern of anomalies including developmental delay, intellectual disability, seizures, dysmorphic facial features, mesomelic dysplasia, and failure to thrive. Although there is some overlap, the clinical presentation of this autosomal dominant disorder appears to be distinct from CHOPS syndrome.

## Material and Methods

### Enrollment

Participants were enrolled after written informed consent was obtained from parents or legal guardians according to ethical review boards policies. The clinical evaluation included medical history interviews, physical examinations and review of medical records. The Deciphering Developmental Disorders (DDD)^19^ identifier of proband 4 is DDD276869.

### Exome/Genome sequencing and analysis

Affected individuals were selected for sequencing to establish a diagnosis.

#### Proband 1

Trio exome analysis was performed on a NextSeq 500 Sequencing System (Illumina, San Diego, CA) after a 12-plex enrichment with SeqCap EZ MedExome kit (Roche, Basel, Switzerland), according to manufacturer’s specifications. Sequence quality was assessed with FastQC 0.11.5, reads were mapped using BWA-MEM (v 0.7.13), sorted and indexed in a bam file (samtools 1.4.1), duplicates were flagged (sambamba 0.6.6), coverage was calculated (picard-tools 2.10.10). Variant calling was done with GATK 3.7 Haplotype Caller. Variants were then annotated with SnpEff 4.3, dbNSFP 2.9.3, gnomAD, ClinVar, HGMD, and an internal database. Coverage for these samples was 93% at a 20x depth threshold.

#### Probands 2 and 10

Exomes were captured using the IDT xGen Exome Research Panel v1.0 for proband 2 and her parents and SureSelect Human All Exon V4 (50 Mb) for proband 10 and his parents. Sequencing and analyses were performed as previously described^20^. The general assertion criteria for variant classification are publicly available on the GeneDx ClinVar submission page.

#### Proband 3

The exomes of proband 3, his parents and two healthy siblings were captured and sequenced as described^21^. Variants were called and filtered using the Varapp software^22^. Sanger sequencing confirmed the anticipated segregation of the potentially causative variants.

#### Proband 4

Exome capture and sequencing was performed as previously described^19^.

#### Proband 5

Exome sequencing of the proband was performed as previously described^23^. Sanger sequencing of samples from parents revealed *de novo* segregation of the variant.

#### Proband 6

Trio genome analysis was performed as previously described^24^. Sanger sequencing confirmed the *de novo* variant reported here.

#### Proband 7

Trio exome analysis was performed as previously described^25^.

#### Proband 8

Sample preparation and enrichment was performed using TruSeq DNA Exome kit (Illumina) and sequencing was performed using NextSeq 500 (Illumina) with mean region coverage 83x. Variant were called using VarAft software. Variant analysis was performed according to standards and guidelines for the interpretation of sequence variants^26^. Sanger sequencing confirmed the *de novo* origin of variant.

#### Proband 9

Trio exome analysis was performed with Agilent SureSelect CRE exome capture, Illumina NextSeq 500 sequencer and a mean coverage of 100x. Data were processed using Cpipe^27^ and variant filtering and prioritization were phenotype driven (gene lists: intellectual disability, Mendeliome). Variant classification followed ACMG guidelines.

### Protein alignment

Alignments of ALF family members were made using Clustal Omega^28^ (v1.2.4) and imported on Jalview^29^ for visualization.

### Interaction modeling

3D modeling for AFF3 (UniProt entry P51826) and SIAH1 (Q8IUQ4) interaction^30^ was obtained on Swiss-PdbViewer-DeepView^31^ v4.1. As no structural model for human SIAH1 ubiquitin-ligase was available, we used mouse ubiquitin ligase structure (pdb 2AN6) 100% conserved with human sequence in the binding region^32^.

### Mouse models

Brain neuroanatomical studies were performed on three 16-week-old male mice in C57BL/6N background with homozygous knock-out of the *Aff3* (a.k.a. *Laf4*) gene^33^. Seventy-eight brain parameters were measured across three coronal sections as described^34^ and data were analyzed using a mixed model and comparing to more than 100 wild-type males using a false discovery rate of 1%. Other metabolic and anatomical phenotypes were assessed by the Welcome Trust Sanger Institute through phenotyping of 6 to 13 homozygous and 7 to 14 heterozygous mice and are available on the International Mouse Phenotyping Consortium website. Engineering of *Aff3*^del^ mice model carrying a 353 kb deletion homologous to the one harbored by an affected individual^15^ was previously published^35^. E18.5 animals were processed and stained as described^36^. With Taconic Biosciences GmbH, Cologne, Germany, we engineered a constitutive *Aff3*^A233T^ knock-in through CRISPR/Cas9-mediated gene editing using TGGTGGATGCACGCCGGTTA as guide (NM_001290814.1, NP_001277743.1). This allowed the insertion of an additional silent mutation that creates an AleI restriction site for analytical purposes.

### Zebrafish overexpression model

Human wild-type ORFs (AFF3, NM_002285.2 and AFF4, NM_014423.4) cloned into the pEZ-M13 vector were transcribed using the mMessage mMachine kit (Ambion) as prescribed. We injected 1-2 nL of diluted RNA (100-300 ng) inside the yolk, below the cell of wild-type zebrafish embryos at the 1- to 2-cell stage. Phenol red dye with distilled water was injected as vehicle control in similar volume. Injected embryos were raised at 28°C and fixed in 4% PFA for 2 hrs at 4-5 days post fertilization (dpf) and stored in PBS at 4°C. Pictures of the embryos were taken after embedding in glycerol. Counts were compared by Fisher exact test.

### Protein accumulation assay

Tagged human wild-type mRNAs cloned into a CMV promoted expression vector were obtained from GeneCopoeia. The ORFs of *AFF3* and *AFF4* were inserted in pEZ-M13 vector with a C-terminal FLAG tag, while the ORF of SIAH1 (NM_001006610) was inserted in pEZ-M07 vector with a C-terminal 3xHA tag. The *AFF3* NM_002285.2:c.697G>A, c.704T>G, and *AFF4* NM_014423.4:c.772C>T mutations were engineered using the QuikChange II XL Site-Directed Mutagenesis Kit (Agilent Technologies) following the manufacturer’s instructions. HEK293T cells cultured in complete medium (DMEM containing 10% FBS and 1% penicillin-streptomycin) were transiently transfected with wild type and mutated plasmids using calcium phosphate. 24 hrs after transfection, medium was changed to fresh complete medium. Total protein extracts were obtained after 48 hrs using RIPA buffer with protease and phosphatase inhibitor cocktail. Denatured protein extracts were immunoblotted with anti-FLAG (F3165), -HA (12CA5) and -β-actin (A2066) antibodies from Sigma-Aldrich.

## Results

We identified ten unrelated affected individuals (probands 1-10) with *de novo* missense variants in the ALF domain of AFF3 (**Figure 1A and Table 1**) through trio-based exome sequencing and data aggregation of multiple laboratories and clinical centers via GeneMatcher^37^. The four different identified variants (**Table 1**) (i) are not present in the Genome Aggregation Database (gnomAD^38^ v2.1.1); (ii) are predicted to be deleterious by SIFT^39^, PROVEAN^40^, PolyPhen2^41^ and MutationTaster2^42^; (iii) are part of the top 1% of all deleterious variants with CADD scores over 20; and (iv) modify highly conserved amino acids (**Figure 1B-C**). Nine of the probands present variants affecting the same codon of exon 6, c.772G>T p.(A258S) (probands 1-2), c.772G>A p.(A258T) (probands 3-8), c.773C>T p.(A258V) (proband 9), whereas proband 10 carries a variant perturbing a neighboring codon c.779T>G p.(V260G) (NM_001025108.1, NP_001020279.1; **Table 1**). An eleventh individual (deletion proband) carrying a 500kb microdeletion and an overlapping phenotype (see below) was previously described^15^. This deletion removes exons 4 to 13 of *AFF3*, which encode its N-terminal region, including the ALF and its degron and part of the transactivation domains and was proposed to act as a dominant negative^35^ (**Figure 1A**).

**Figure 1:**
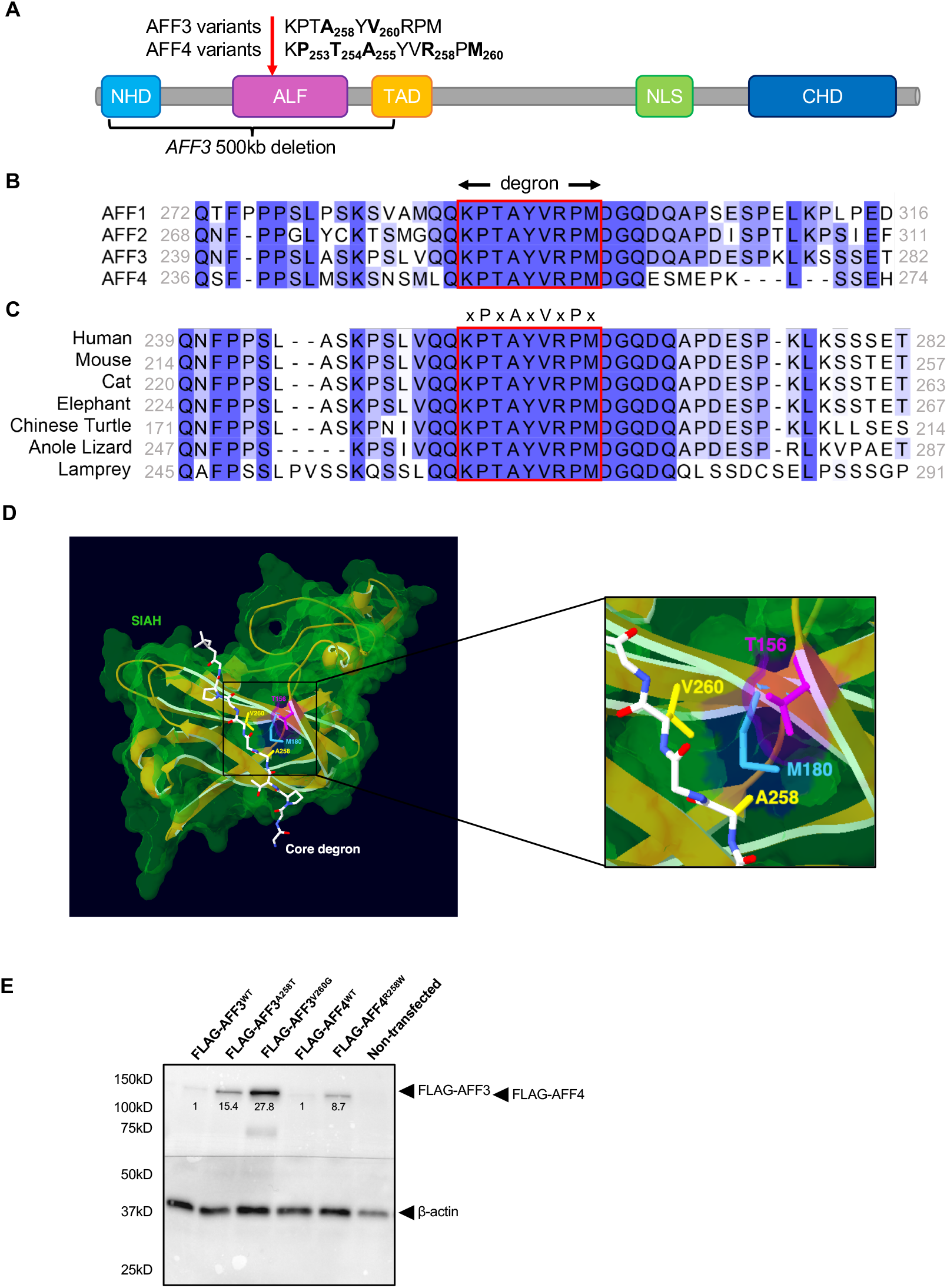
AFF3 and AFF4 degron motif variants. **(A)** Schematic protein structure of ALF proteins with from the amino terminus a N-terminal homology domain (NHD), the AF4/LAF4/FMR2 homology domain (ALF)^21^ containing the SIAH-binding degron motif, a serine-rich transactivation domain (TAD)^3^, a bipartite nuclear/nucleolar localization sequence (NLS), and a C-terminal homology domain (CHD). The sequences of the degron motif of AFF3 and AFF4 are shown above. The residues modified in the KINSSHIP probands described in this manuscript and individuals affected by CHOPS^16, 17^ are highlighted in bold and numbered. The extent of the 500kb deletion previously described^35^ is indicated here. A red arrow pinpoints the position of the degron motif. **(B)** Amino acid sequences alignment of human AFF1, AFF2, AFF3 and AFF4 proteins (ENSP00000305689, ENSP00000359489, ENSP00000317421 and ENSP00000265343, respectively) showing the highly conserved degron motif (red rectangle) of the ALF homology domain that provides the binding moiety to the SIAH ubiquitin-ligase. Sequences alignment was performed using Clustal Omega and edited using Jalview. Shading is proportional to conservation among sequences. **(C)** Amino acid sequences alignment of different AFF3 vertebrate orthologs showing the conservation of the degron motif (red rectangle). Accession numbers are ENSP00000317421 (human), ENSMUSP00000092637 (mouse), ENSFCAP00000024603 (cat), ENSLAFP00000010776 (elephant), ENSPSIP00000007060 (chinese turtle), ENSACAP00000008035 (anole lizard) and ENSPMAP00000008605 (lamprey). **(D)** 3D modeling of the binding of human AFF3 degron to the mouse Siah ubiquitin ligase. PDB entry 2AN6^32^ was loaded in Swiss-PdbViewer^31^ and used as a template to align the human SIAH ubiquitin ligase (uniprot entry Q8IUQ4)^30^. With respect to the mouse crystal structure, the only difference is the presence of an aspartic acid residue instead of a glutamic acid at position 116. The region of AFF3 containing the degron motif (LRPVAMVRPTV) was then aligned onto the Siah-interacting protein^50^ peptide present in the crystal structure (QKPTAYVRPMD) to highlight the position of the variants reported in this study. For clarity, only side-chains of the core degron motif (P256, A258, V260 and P262) are shown, with yellow highlights on the KINSSHIP mutated residues. A zoom in is displayed on the right. The core degron motif adopts a beta-strand conformation directly contacting the ubiquitin ligase-binding groove. The sidechains of A258 and V260 are embedded into binding pockets too small to accommodate larger side chains^32^. They are in direct proximity of Siah residues T156 (pink) and M180 (cyan), identified as key binding residues in a series of pull-down assays^32^. The longer side-chains of A258T, A258S, A258V variants and the smaller V260G are likely to weaken or prevent the interaction with the ligase. **(E)** Immunoblot showing the accumulation of mutated forms of AFF3 and AFF4 proteins compared to wild type (WT). Protein extracts of HEK293T cells independently expressing FLAG-tagged AFF3^WT^, AFF3^A258S^, AFF3^V260G^, AFF4^WT^ and AFF3^R258W^ proteins were immunoblotted with an anti-FLAG antibody (upper portion) and an anti-ß-actin antibody for loading control (bottom portion). The positions of FLAG-AFF3, FLAG-AFF4 and ß-actin proteins are indicated on the right; they are 133kD, 127kD and 42kD respectively. Signal intensity is measured and normalized on corresponding loading control.

**Table 1.** Predicted pathogenicity and allele frequencies of *AFF3 de novo* variants

| Gene | Individual | Chromosome coordinates (GRCh37/hg19) | Nucleotide change | Amino acid change | dbSNP (v152) | GnomAD allele frequency (v2.1.1) | Deleteriousness prediction (score) |  |  |  |  |
| --- | --- | --- | --- | --- | --- | --- | --- | --- | --- | --- | --- |
|  |  |  |  |  |  |  | CADD_PHRED (GRCh37-v1.4) | SIFT (v4.0.3) | PROVEAN (v1.1) | PolyPhen2 (v2.2.2) | Mutation Taster2 (09.01.19) |
| *AFF3*<br>NM_001025108.1<br>NP_001020279.1 | 1 | Chr2:100623270 | c.772G>T | A258S | - | 0 | 23.8 | Damaging (0.000) | Neutral (-2.48) | Probably damaging (1.000) | Disease causing |
|  | 2 |  |  |  |  |  |  |  |  |  |  |
|  | 3 | Chr2:100623270 | c. 772G>A | A258T | rs1131692272 | 0 | 24.3 | Damaging (0.000) | Deleterious (-3.30) | Probably damaging (1.000) | Disease causing |
|  | 4 |  |  |  |  |  |  |  |  |  |  |
|  | 5 |  |  |  |  |  |  |  |  |  |  |
|  | 6 |  |  |  |  |  |  |  |  |  |  |
|  | 7 |  |  |  |  |  |  |  |  |  |  |
|  | 8 |  |  |  |  |  |  |  |  |  |  |
|  | 9 | Chr2:100623269 | c.773C>T | A258V | - | 0 | 24.2 | Damaging (0.000) | Deleterious (-3.30) | Probably damaging (1.000) | Disease causing |
|  | 10 | Chr2:100623263 | c. 779T>G | V260G | - | 0 | 24.4 | Damaging (0.000) | Deleterious (-5.85) | Probably damaging (1.000) | Disease causing |

**Footnote:** SIFT cutoff=0.05, PROVEAN cutoff=-2.5, PolyPhen2 cutoff=0.5.

All *AFF3* variants described here and CHOPS syndrome-associated *AFF4 de novo* missense previously published^16, 17^ map within the degron motif of the ALF domain. This highly conserved 9 amino acid sequence [xPxAxVxPx] (**Figure 1A-B**) mediates interaction with the SIAH E3 ubiquitin ligase and regulates their degradation^1^. According to pathogenic variant enriched regions (PER)^43^, the degron is predicted to be constrained within the ALF family. Pathogenicity of the four *de novo AFF3* identified variants is further supported by the three-dimensional representation of part of the encoded peptide (**Figure 1D**). The mutated residues are located within the degron motif (KPT**A_258_**Y**V_260_**RPM), which adopts a beta-strand conformation directly contacting the SIAH ubiquitin ligase binding groove^30^. The side chains of Alanine 258 and Valine 260 are embedded into the hydrophobic core of the beta-sandwich where the binding pockets are too small to accommodate larger side chains^32^. Thus, the variants p.(A258T), p.(A258S), p.(A258V) and p.(V260G) are likely to weaken or prevent binding to the ubiquitin ligase. Hence, all these *de novo* variants, as well as the 500kb deletion previously reported^15^ that encompasses the degron, could result in hindered degradation and thus accumulation of AFF3. Consistent with this hypothesis, transiently transfected FLAG-tagged AFF3^A258S^ and AFF3^V260G^ proteins were more stable than wild-type FLAG-tagged AFF3 (**Figure 1E**). The previously reported AFF4 *de novo* variants p.(P253R), p.(T254A), p.(T254S), p.(A255T), p.(R258W) and p.(M260T) that also affect the degron motif (K**P_253_T_254_A_255_**YV**R_258_**P**M_260_**) (**Figure 1A**) were similarly shown to reduce clearance of the ALF transcription factor by SIAH^16, 17^.

We compared the phenotypes of the ten individuals with *de novo* variants in *AFF3* described here and that of the previously reported case carrying *AFF3* partial deletion^15^ (**Table S1** for detailed phenotypes). They exhibit severe developmental epileptic encephalopathy (10 probands out of 11), along with mesomelic dysplasia resembling Nievergelt/Savarirayan mesomelic skeletal dysplasia (NSMSD) (10/11) and failure to thrive (10/11). These three features are often associated with microcephaly (7/11), global brain atrophy and/or ventriculomegaly (7/9) (**Figure S1**), fibular hypoplasia (9/11), horseshoe kidney (8/11), abnormalities of muscle tone (9/10), gastroesophageal reflux disease (5/10) and other gastrointestinal symptoms (10/10). They also share common dysmorphic facial features such as a bulbous nasal tip (6/9), a wide mouth with square upper lip (7/10), abnormalities of the teeth and gums (9/10) and hypertrichosis (8/9) (**Figure 2–3**). Respiratory difficulties/pulmonary involvement were observed in about half of the probands with *de novo* variants (6/11). Whereas respiratory arrest led to the death of proband 3 at 21 years, the deletion proband died at four months after recurrent apneic episodes (**Table S1**).

**Figure 2:**
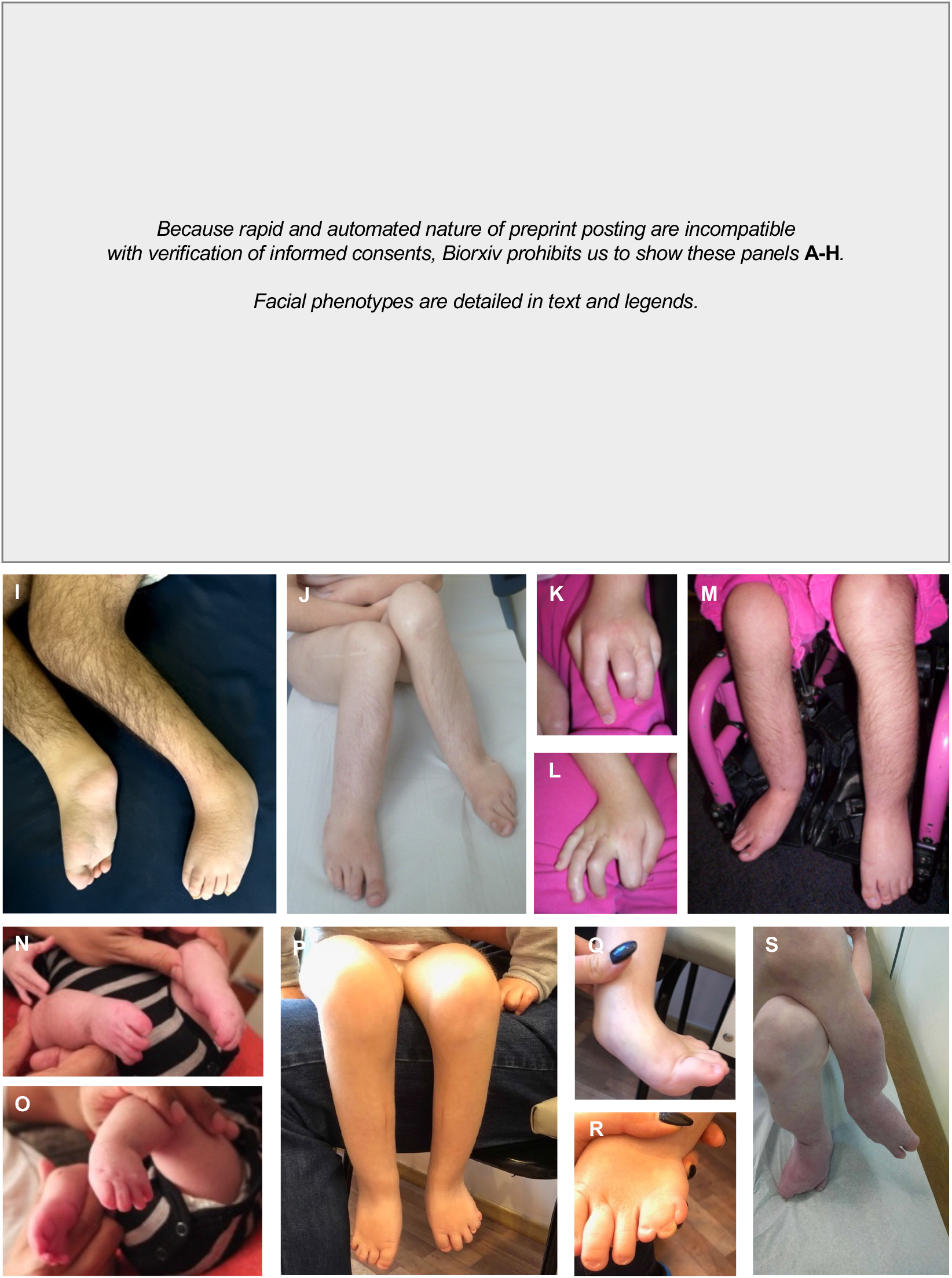
Photographs of KINSSHIP individuals with *AFF3 de novo* missense variants. **(A)** Proband 2 at 2 years 6 months old; **(B, I)** Proband 3 at 18 years old; **(C)** Proband 4 at 9 months and **(D, J)** 21 years old; **(E)** Proband 5 at 1 year 7 months and **(N-O)** 16 days old; **(F, K-M)** Proband 6 at 9 years old; **(G)** Proband 7 at 8 years old; **(H, P-R)** Proband 8 at 7 years 9 months old; **(S)** Proband 10 at 11 years old. Note the synophrys and micrognathia, protruding ears, large nose with prominent nasal tip and prominent teeth in proband 3 **(B)**, 4 **(D)**, 6 **(F)** and 8 **(H)**, as well as their hypertrichosis of the limbs **(I, J, M, P)**. Together with probands 5 and 7, they exhibit thick hair, long eyelashes and a wide mouth **(E, G)**. Facial features coarsen with age as shown by pictures of proband 4 at different ages **(C-D)**, explaining the more delicate features of younger probands **(A, E)**. *AFF3 de novo* missense variant carriers also have hypoplastic talipes and abnormalities of toes **(I, J, M-S)**. Proband 6 also shows clinodactyly and soft tissue syndactyly of both hands **(K, L)**.

**Figure 3:**
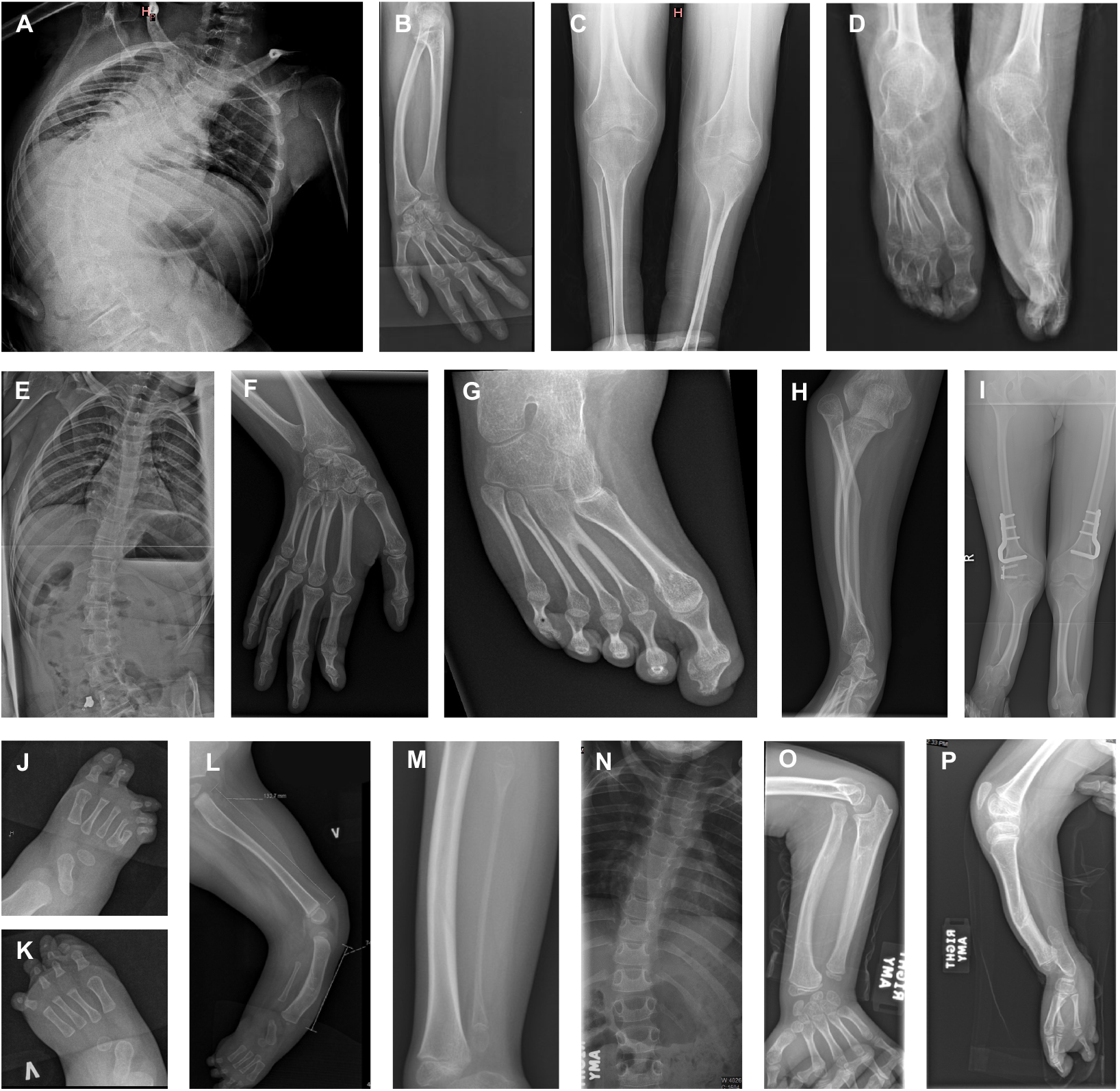
X-rays of KINSSHIP individuals with *de novo* missense variants in *AFF3*. **(A-D)** Proband 3 at 18 years old; **(E-I)** Proband 4 at 21 years old; **(J-L)** Proband 5 at 10 months old; **(M)** Proband 7 at 8 years old; **(N-P)** Proband 10 at 10 years old. **(A)** Severe scoliosis and fusion of C2-C3 vertebral bodies and L5-S1 vertebral cleft **(B)** Dorsal and radial bowing of the radius and “V-shaped” proximal carpal bones as seen in Madelung deformity, **(C)** metaphyseal widening and hypoplastic fibula and **(D)** hypoplastic talipes. **(E)** Static scoliosis, **(F)** Short ulna and radius and bilateral dislocation/subluxation of radial heads. Note erratic articulation of the styloid process of the ulna on the radius rather than on the carpal bones, **(G)** Congenital fusion of the bases of the second and third right metatarsals, **(H)** Hypoplastic bowing femora and **(I)** short tibias with enlarged metaphyses. **(J)** Right foot with 4^th^ and 5^th^ metatarsals synostosis **(K)** and left foot missing the lateral ray, **(L)** Extremely short rectangular fibula and bowed tibia. **(M)** Hypoplastic fibula. **(N)** Scoliosis and cervical ribs, **(O)** bowed radius and ulna, **(P)** bowed tibia, severely hypoplastic fibula and oligodactyly.

This constellation of features recalls some features of CHOPS-affected individuals. The three originally described probands^16^, along with the eight recently identified^17^, presented with distinctive facial dysmorphic features reminiscent of CdLS, short stature with obesity (11/11), developmental delay/intellectual disability (DD/ID) (11/11) and microcephaly (6/11) without epilepsy. They showed gastrointestinal abnormalities (8/11), accompanied by abnormal feeding behavior (6/6), hearing loss (8/11), cardiac (8/11) and pulmonary defects (8/11) and rarely horseshoe kidney (2/11). Whereas they present with vertebral abnormalities (5/11) and brachydactyly (8/11), mesomelic dysplasia is never observed and hypoplastic fibula rarely (1/11).

Although phenotypes of *AFF3* and *AFF4* missense carriers are overlapping, they are not identical. We thus suggest naming the distinct autosomal dominant AFF3-associated disorder KINSSHIP syndrome (<u>KI</u>dney anomalies, <u>N</u>ievergelt/<u>S</u>avarirayan mesomelic dysplasia, <u>S</u>eizures, <u>H</u>ypertrichosis and <u>I</u>ntellectual disability with <u>P</u>ulmonary involvement, MIM #XXXX) to evoke both some of its cardinal characteristics, as well as its similarity (common mode of action and inheritance and overlapping phenotypes) with CHOPS syndrome.

To better understand the functional effects of *AFF3* variation, we investigated both knock-out and knock-in mouse models (**Table 2**). We first studied the knock-out mouse line engineered by the International Mouse Phenotyping Consortium^33^ (IMPC). The IMPC routinely measures an extensive series of parameters and evaluate if those are significantly different from wild-type mice^44^ (*p*≤10^−04^). *Aff3*^+/-^ and *Aff3*^−/−^ mice exhibit skeletal defects including fusion of vertebral arches, vertebral transformation and decreased caudal vertebrae number. Homozygous knock-out mice also show an abnormal skull shape with a small, deviated snout and malocclusion as well as decreased serum fructosamine and albumin levels that could reflect kidney defects and/or metabolic dysregulation. Neurological dysfunctions were also noted with an increased or absent threshold for auditory brainstem response (signs of hearing impairment) and diminished grip strength. As *Aff3* is expressed in progenitor neurons^45^ and required for neuronal migration in the cerebral cortex^46^, we further assessed the consequences of *Aff3* disruption on brain development by measuring a standardized set of 78 parameters across 22 brain regions^34^. Compared with wild type males, homozygous *Aff3*^−/−^, but not heterozygous *Aff3*^+/-^ males, exhibited significantly enlarged lateral ventricles (*p* = 1.24 × 10^−04^) and decreased corpus callosum size (*p* = 3.02 × 10^−06^; **Figure 4**), similar to the phenotypes observed in proband 2, 3 and 6 and in the previously reported deletion proband (**Table S1**, **Figure S1**)^15^. These features are in stark contrast with results obtained with another engineered *Aff3*^−/−^ line that showed no phenotypic perturbations possibly because of genetic background differences, i.e. C57BL/6N versus CD1, and/or focusing on limb morphology only^35^.

**Table 2.** AFF3 mice models and their phenotypes

**Footnote:** SIFT cutoff=0.05, PROVEAN cutoff=-2.5, PolyPhen2 cutoff=0.5.
| Table 2: AFF3 mice models and their phenotypes |  |  |  |  |  |  |  |
| --- | --- | --- | --- | --- | --- | --- | --- |
| Producer |  | IMPC<sup>33</sup> |  | Kraft et. al.<sup>35</sup> | This work |  |  |
| Background |  | *C57BL/6N* |  | CD1 | CD1 |  | *C57BL/6N* |
| Genotype |  | *Aff3*<sup>+/-</sup> | *Aff3*<sup>-/-</sup> | *Aff3*<sup>-/-</sup> | *Aff3*<sup>del/+</sup> | *Aff3*<sup>del/del</sup> | *Aff3*<sup>A233T</sup> |
| Phenotype | Craniofacial anomalies | - | + | NA | Variable features, from non-affected to homozygous-like phenotype | + | NA |
|  | Vertebral malformations | + | + | NA |  | NA |  |
|  | Mesomelic dysplasia | - | - | - |  | + * |  |
|  | Polydactyly | - | - | - |  | + |  |
|  | Kidney malfunction | - | + | NA |  | NA |  |
|  | Intestinal prolapse | - | - | NA |  | + |  |
|  | Neurological dysfunctions | - | + | NA |  | + |  |
|  | Neuroanatomical defects | NA | + | NA |  | NA |  |
|  | Reduced body size | - | - | NA |  | + |  |
| Lethality |  | - | - | NA | Low chimerism, no heterozygous offspring, suggesting lethality | Postnatal lethality | No heterozygous offspring, suggesting lethality |
| NA: Not available; IMPC significance threshold $p \leq 10^{-04}$, *Kraft et. al<sup>35</sup> | | | | | | | |

**Figure 4:**
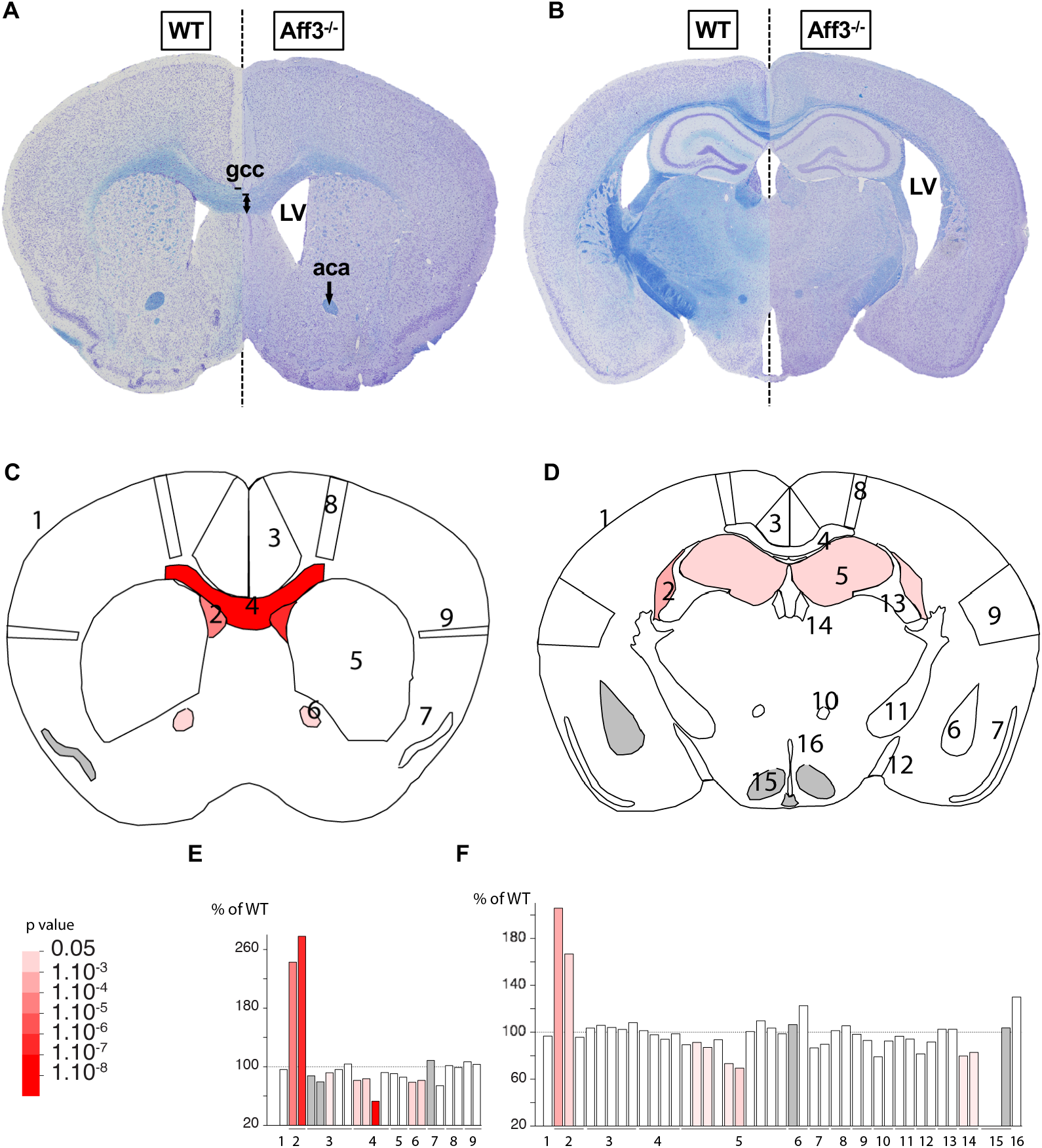
Neuroanatomical defects in *Aff3*^−/−^ mice. Merged double-stained sections in *Aff3*^−/−^ mice (right of dashed lines) and their matched controls (WT: wild type, left of dashed lines) at the striatum **(A)** and at the hippocampus **(B)** levels with schematic representation of the affected areas **(C, D)**. Histograms showing the percentage of increase or decrease of parameters in measured areas as compared to the controls, for striatum **(E)** and hippocampus **(F)** sections. Red shading is proportional to the stringency of the significance threshold. Numbers indicate studied areas: 1=total brain area, 2=lateral ventricles, 3=cingulate cortex (section 1) and retrosplenial cortex (section 2), 4=corpus callosum, 5=caudate putamen (section 1) and hippocampus (section 2), 6=anterior commissure (section 1) and amygdala (section 2), 7=piriform cortex, 8=motor cortex, 9=somatosensory cortex, 10=mammilo-thalamic tract, 11=internal capsule, 12=optic tract, 13=fimbria, 14=habenular, 15=hypothalamus, and 16=third ventricle. Results demonstrate an enlargement of lateral ventricles (**LV**; p=1.24E-04 on section 1, p=4.64E-02 on section 2) and a smaller genu of the corpus callosum (**gcc**; decreased corpus callosum size p=6.35E-02 indicated by the black dash and double arrow, decreased bottom width of the corpus callosum p=3.02E-06 and decreased height of the corpus callosum p=4.96E-02). Other phenotypes of lesser stringency can be observed such as atrophy of the anterior commissure (**aca**; p=1.02E-02) and smaller hippocampus (p=4.02E-02).

We then reassessed mouse models mimicking the deletion identified in the previously described proband, which were previously engineered to assess an aggregation method for the rapid generation of structural variants^35^. Consistent with the phenotype of the deletion proband, homozygous animals chimeric for a 353kb deletion syntenic to the 500kb human deletion exhibited mesomelic dysplasia, triangular tibia, severe hypoplastic fibula and polydactyly of the feet^35^ (**Table 2**). Reexamination of these *Aff3*^del/del^ (a.k.a. *Laf4^del/del^*) mice showed that they also presented with reduced body size, craniofacial dysmorphisms with delayed ossification of skull bones, hypoplastic pelvis, intestinal prolapse and neurological dysfunction (**Figure 5A-C**). Chimeric *Aff3*^del/+^ heterozygotes presented with variable features ranging from unaffected to homozygous deletion-like phenotypes. Whereas *Aff3*^del/+^ animals with low chimerism were fertile they produced no heterozygous offspring suggesting lethality of the 353kb deletion (**Table 2**). While these results support a causative role for the deletion in the deletion proband, they do not allow differentiating between gain-of-function and haploinsufficiency.

**Figure 5:**
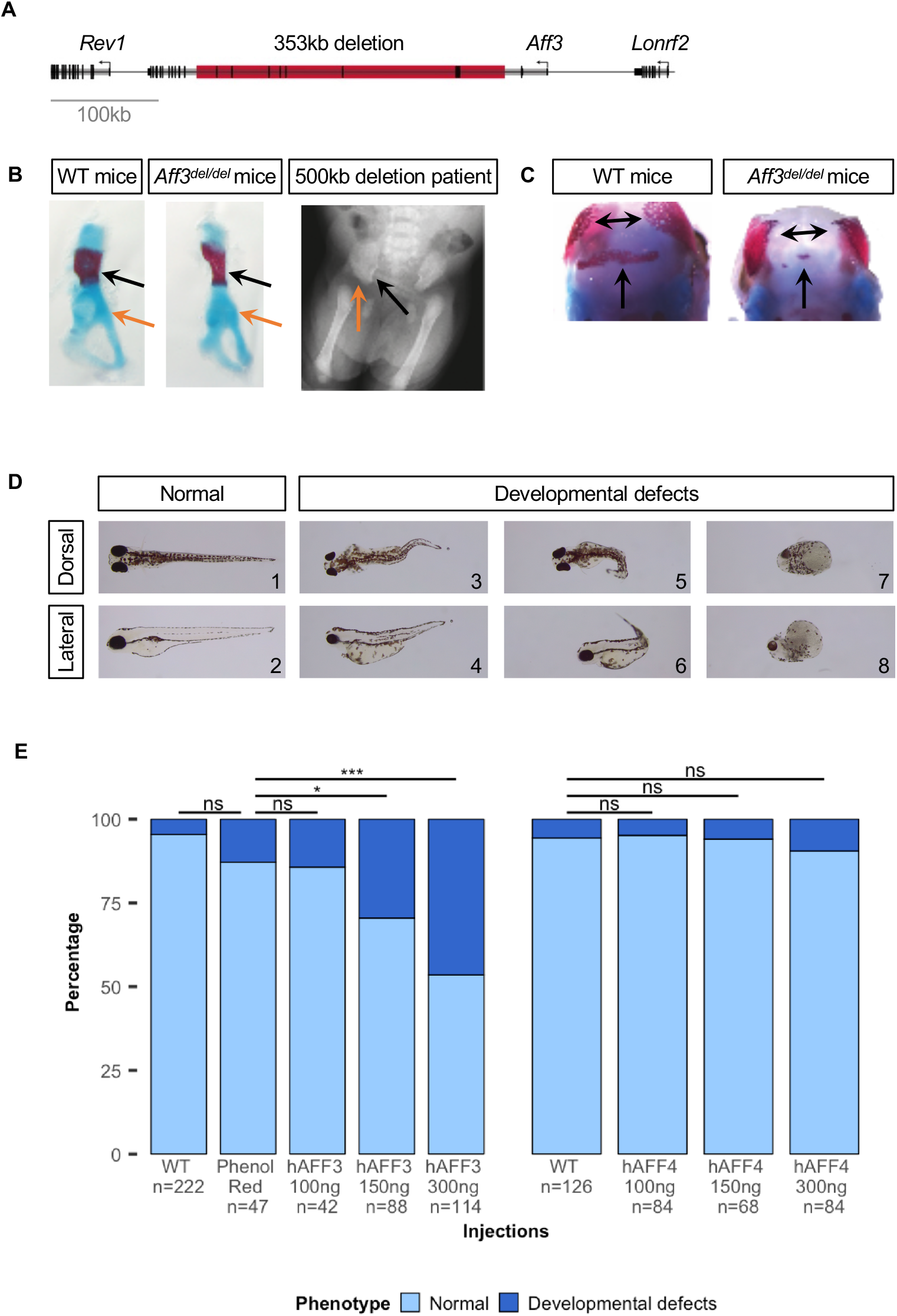
Animal models. **(A)** Schematic representation of the deletion generated in mice ES cells with the CRISPR/Cas9 system, which models the mutation observed in the deletion proband^15, 35^. **(B)** Skeletal staining of E18.5 mouse embryos show mesomelic dysplasia with triangular tibia and hypoplastic fibula (see Figure 3 in reference^35^), as well as a hypoplastic pelvis in *Aff3^del/del^* mice, especially noticeable in the iliac wing (black arrows) and acetabulum (orange arrows), perturbations also observed in the deletion proband. **(C)** Delayed ossification of flat bones in the skull of *Aff3^del/del^* mice. **(D)** Lateral (top line) and dorsal (bottom line) views of the observed phenotypes of 4 dpf AB-WT zebrafish embryos injected with human AFF3 mRNA (hAFF3). hAFF3-injected zebrafish embryos exhibit severe developmental defects including a bent body axis and yolk sac edema (D3-6), as well as extreme malformations with absence of body axis, tail and fins and cyclopia (D7-8). Embryos with normal development are displayed for comparison (D1-2). **(E)** Proportions of normal and developmentally defective 4 dpf AB-WT zebrafish embryos upon injection of increasing doses of hAFF3 (left panel) and hAFF4 (right panel) mRNA. Dark and light colors indicate developmentally defective and normal animals, respectively. Control injections with Phenol Red show no significant (ns) differences with WT in both AFF3 and AFF4 experiments (Fisher exact test, p=0.09 and p=0.12 respectively). hAFF3 mRNA injection significantly increases the number of zebrafish with developmental defects when compared to controls starting from 150ng (*; p=0.03) and reinforced at 300ng (***; p=3.2E-5). AFF4 injections do not have a significant impact on zebrafish development compared to WT, even at the same dose (300ng, p=0.29).

To further assess the underlying mutational mechanism in our missense probands, we engineered a knock-in mouse model carrying the *Aff3*^A233T^ mutation that is the equivalent of the most commonly observed *de novo* variant, identified in probands 3 to 8 [p.(A258T)]. The microinjection of a total of 410 C57BL/6NTac zygotes and transfers into 14 recipient females to allow CRISPR/Cas9 editing resulted in only 13 pups at weaning. Genotyping showed that most of them were either wild type (8 individuals) or carried CRISPR/Cas9-mediated mutations (4) although reduced gRNA activity was used for microinjection. A single female F0 founder animal showed the targeted A233T knock-in but with a very low mosaicism rate of 16.7% in an ear biopsy. Genotyping showed that none of its offspring from four consecutive pregnancies were heterozygous for the mutation. These results suggest that the *Aff3*^A233T^ mutation is lethal with high mosaicism (homozygous *Aff3*^A233T/A233T^ and heterozygous *Aff3*^+/A233T^ chimeras), in gametes or during the fetal period (heterozygous *Aff3*^+/A233T^; **Table 2**). The success statistics of similar CRISPR/Cas9 knock-in projects performed by Taconic Biosciences GmbH (Cologne, Germany) through the years further support this hypothesis. Out of 92 attempted knock-in constructs 98% were successful with only 2% failing to generate F0 animals. For most projects, positive F1 animals were also generated.

To lend further support to the model centered on a pathological increase of AFF3 protein product in affected individuals, we assessed its accumulation in zebrafish. Whereas the genome of these teleosts encodes four ALF transcription factors orthologous to the mammalian *AFF1* to *AFF4*, these genes do not harbor a [xPxAxVxPx] degron motif suggesting that their degradation is regulated differently in fish. Therefore, we modeled accumulation by independently overexpressing increasing amounts of unmutated human AFF3 and AFF4 mRNA in zebrafish embryos. We observed a dose-dependent increase in the fraction of 4 dpf embryos with morphological defects upon overexpression of AFF3. The observed phenotypes included bent body axis, yolk sac edema and generalized body development defects at higher doses (**Figure 5D-E**). A similar albeit less pronounced dose-dependent increase in zebrafish embryos with morphological defects was seen upon overexpression of AFF4 (**Figure 5E**).

To further assess the redundancy of ALF transcription factors, we took advantage of published knockdown experiments^47^. Luo and colleagues established and profiled the transcriptome of stable HEK293T cell lines independently knocked down for *AFF2, AFF3* and *AFF4* expression by specific shRNAs. Reanalysis of these data confirmed that ALF transcription factors have mostly different target genes, as 55% (125 out of 226), 62% (261/423) and 87% (966/1116) of the genes are specifically perturbed by the knock down of *AFF2, AFF3* and *AFF4*, respectively (**Figure 6A**). Intriguingly, the subset of common targets is similarly influenced by decreased expression of *AFF2* and *AFF3* (**Figure 6C, D**), whereas knocking down *AFF3* and *AFF4* had opposite effect (**Figure 6B, C**). 95% (119 out of 125) of common targets are decreased upon reduction of *AFF3* and increased upon reduction of *AFF4* expression suggesting that these two transcription factors act as positive and negative regulators of common pathways. Within the genes perturbed by both *AFF3* and *AFF4*, we observed a significant overrepresentation of genes implicated in the gastrin hormone pathway (CCKR signaling map, P06959) and a proton pump complex (vacuolar proton-transporting V-type ATPase complex, GO:0016471) possibly associated with the gastroesophageal reflux disease observed in both KINSSHIP and CHOPS individuals. Genes linked to the gonadotropin-releasing hormone receptor pathway are similarly enriched (P06664). This observation could be related to cryptorchidism of KINSSHIP proband 1 and small genitalia/cryptorchidism in three out of five males affected by CHOPS syndrome^16, 17^, as well as the erratic menstrual cycle of proband 4 (most probands being too young to predict any pubertal anomaly) and popliteal pterygium in proband 8 (**Table S1**).

**Figure 6:**
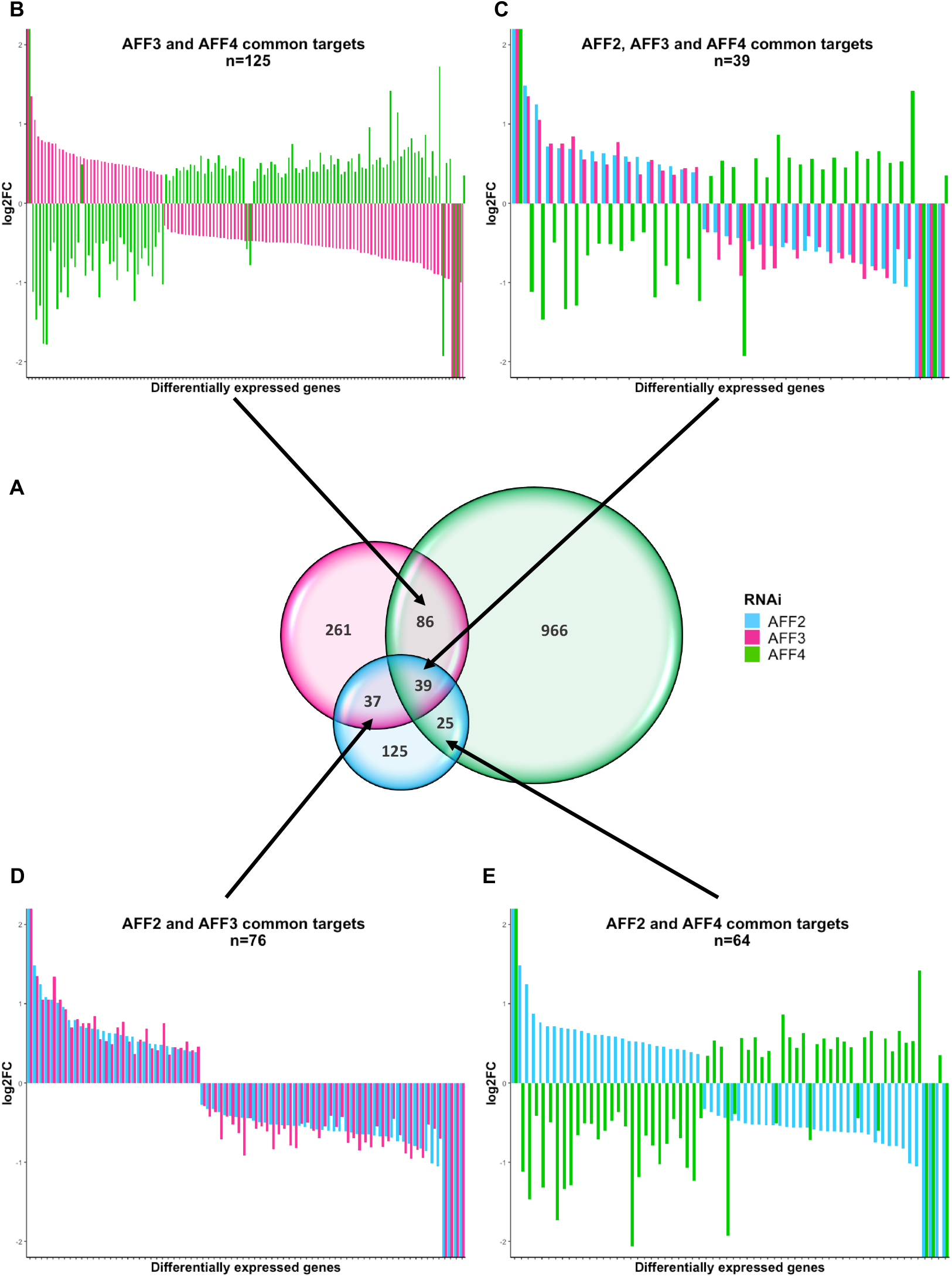
*AFF2, AFF3* and *AFF4* targets. **(A)** Venn diagram of the differentially expressed genes from independent RNAi of *AFF2, AFF3*, and *AFF4* (adapted from Luo et al^47^) showing that ALF transcription factors have different targets. The knocked-down gene color code is indicated on the right. The expression modifications of common targets are presented in panels **B-E**. Common differentially expressed genes show adverse regulation between knockdown of *AFF3* and *AFF4* **(B)**, *AFF2* and *AFF4* **(E)** and *AFF2/AFF3* and *AFF4* **(C)**. Whereas RNAi of *AFF2* and *AFF3* similarly influence their 76 common targets **(D)**, knockdowns of *AFF2* or *AFF3* have opposite effects to that of *AFF4* on their common targets [54/64=84% **(E)** and 119/125=95% **(B)** of common targets with opposite perturbation of their expression levels, respectively].

## Discussion

All eleven individuals with an *AFF3* variant we identified have a complex but overlapping clinical presentation, which we named KINSSHIP syndrome. One of the cardinal characteristics of this rare autosomal dominant syndrome is mesomelic dysplasia with short forearms, radial head dislocation/subluxation, triangular and/or short tibia, fibular hemimelia, hip dislocation, tarsal and/or metatarsal synostosis resembling NSMSD (**Figure 3**). NSMSD is a sporadic or rare autosomal dominant condition^48, 49^ associated with neurodevelopmental and often urogenital abnormalities^50, 51^. KINSSHIP affected individuals similarly present with vertebral and bone mineralization defects, scoliosis, epilepsy, severe global DD/ID sometimes associated with structural brain abnormalities, significant feeding difficulties, horseshoe kidney, hypertrichosis and recognizable facial features. Multiple probands showed coarsening facial features with age, including a large nose with bulbous nasal tip, a prominent columella and a wide mouth with square upper lip (**Figure 2–3, Table S1, Figure S1**).

*AFF3* is one of the targets of the Wnt/β-catenin pathway, an important contributor to pathways involved in bone development and homeostasis^52, 53^. Variants in *WNT* genes cause a diverse range of skeletal dysplasias including mesomelic defects (*WNT5A;* Robinow syndrome, dominant type, OMIM#180700), decreased bone density (*WNT1;* Osteogenesis imperfecta, type XV, OMIM#615220) and limb hypoplasia–reduction defects including fibular a/hypoplasia (*WNT3* and *WNT7A*; Tetra-amelia OMIM#273395 and Fuhrmann syndrome OMIM#228930, respectively). Of note individuals with Robinow rhizo/mesomelic dysplasia also present with developmental kidney abnormalities^54^, whereas perturbations of the Wnt/b-catenin pathway have been associated with the development of ectodermal appendages like hair and teeth^55^. Nine out of ten KINSSHIP probands show dental anomalies. While widespread hypertrichosis may have been partially caused by multi-drug, antiepileptic treatment in probands 3 and 4, its presence in the only non-epileptic *AFF3* individual (proband 9) and the much younger proband 5 seems to confirm the association of this feature with *AFF3* genetic variants. It is possible that the complex clinical presentation of the cases described here (**Table S1**) may represent the effects of impaired *AFF3* function on a number of downstream targets within the Wnt/β-catenin pathway. In-depth transcriptome analysis of affected individuals and/or animal models is warranted to confirm this hypothesis.

Despite the limited number of individuals for both conditions, similarities and differences are notable between individuals with KINSSHIP and CHOPS^16, 17^ syndrome. Individuals with variants in *AFF3* and *AFF4* share features that include respiratory difficulties and vertebral abnormalities, as well as less specific clinical findings such as microcephaly, DD/ID and gastroesophageal reflux disease. Although skeletal abnormalities are reported in both CHOPS and KINSSHIP syndrome, KINSSHIP individuals present with mesomelic dysplasia, whereas CHOPS individuals show brachydactyly. Seizures and failure to thrive are specific to KINSSHIP and obesity with short stature to CHOPS. Congenital heart defects and hearing loss are typically observed in CHOPS, while horseshoe kidney and hypoplastic fibula are predominantly present in KINSSHIP (80% versus 18% and 80% versus 9% of affected individuals, respectively). Despite having thick hair and coarse facies in common, CHOPS probands differ from KINSSHIP probands by their round face and dysmorphic features resembling those of CdLS individuals^16, 17^.

Although proteins encoded by *AFF2*, *AFF3* and *AFF4* were reported to be functionally redundant, at least in regulating splicing and transcription during normal brain development^56^, the clinically distinct phenotypes of individuals carrying *de novo* variants in the degron of AFF3 and AFF4 and our zebrafish results suggest that the encoded proteins are not fully redundant. Further support for this hypothesis is provided by the intolerance to LoF variant of *AFF1* (pLI=0.8), *AFF2* (pLI=1), *AFF3* (pLI=1) and *AFF4* (pLI=1) reported by GnomAD. Whereas homozygous *Aff3*^−/−^ knockout mice display features comparable to those presented by KINSSHIP individuals such as skeletal anomalies, kidney defects, brain malformations and neurological anomalies, these animals do not recapitulate the characteristic mesomelia contrary to *Aff3*^del/del^ mice model. This result and the aforementioned intolerance to LoF suggest that *AFF3* could be associated with two different syndromes, the one described here caused by missense degron variants and a hemizygous deletion of the degron, as well as a second one associated with LoF variants for which affected humans remain to be identified. Although this hypothesis warrants further investigation, we have identified by exome sequencing an individual with features partially overlapping those of KINSSHIP. He is compound heterozygote for a truncating mutation and a predicted to be deleterious missense variant outside of the degron.

In conclusion we describe a new pathology that we propose to name KINSSHIP syndrome. It is associated with variants in the degron of *AFF3* that affect the degradation of the encoded protein. This syndrome shows similarities with the *AFF4*-associated CHOPS syndrome, in particular its gain of protein stability and affected tissues. However, specific KINSSHIP features such as mesomelic dysplasia combined with horseshoe kidney allow a differential diagnosis.

## Supplemental Data

Supplemental data include 1 figure and 1 table.

Figure S1: Brain MRI of proband 7 carrying a de novo variant in AFF3

Table S1: Phenotype of individuals with AFF3 variants

## Acknowledgments

We thank the probands and their families for their participation in this study. We are grateful to Jacques S. Beckmann, Giedre Grigelioniene and the Genomic Technologies Facility of the University of Lausanne. This work was supported by grants from the Swiss National Science Foundation (31003A_182632 to AR), the Simons Foundation (SFARI274424 to AR and SFARI337701 to WKC) and NHGRI (grant UM1HG007301 to SMH, EMB, GMC). The DDD study presents independent research commissioned by the Health Innovation Challenge Fund [grant number HICF-1009-003], a parallel funding partnership between Wellcome and the Department of Health, and the Wellcome Sanger Institute [grant number WT098051]. The views expressed in this publication are those of the author(s) and not necessarily those of Wellcome or the Department of Health. The study has UK Research Ethics Committee approval (10/H0305/83, granted by the Cambridge South REC, and GEN/284/12 granted by the Republic of Ireland REC). The research team acknowledges the support of the National Institute for Health Research, through the Comprehensive Clinical Research Network. This study makes use of DECIPHER which is funded by the Wellcome. We acknowledge the Sanger Mouse Genetics Project for providing mouse samples, funded by the Wellcome Trust grant number 098051. The Australian National Health & Medical Research Council (NHMRC) provided funding for sequencing proband 9 under the Australian Genomics Health Alliance (GNT1113531); the contents are solely the responsibility of the individual authors and do not reflect the views of the NHMRC. The research conducted at the Murdoch Children’s Research Institute was supported by the Victorian Government’s Operational Infrastructure Support Program. The funders had no role in study design, data collection and analysis, decision to publish, or preparation of the manuscript.

## Web Resources

ClustalOmega: http://www.clustal.org/omega/

DDD: http://www.ddduk.org/

GeneMatcher: https://genematcher.org/

GeneDx ClinVar submission page: http://www.ncbi.nlm.nih.gov/clinvar/submitters/26957/

GnomAD: https://gnomad.broadinstitute.org/about.

IMPC: http://www.mousephenotype.org/

MutationTaster2: http://www.mutationtaster.org/

PANTHER: http://www.pantherdb.org

PER viewer: http://per.broadinstitute.org/

PolyPhen-2: http://genetics.bwh.harvard.edu/pph2/index.shtml

PROVEAN: http://provean.jcvi.org/index.php

SIFT: http://sift.jcvi.org/

Varaft: https://varaft.eu

Varapp: https://varapp-demo.vital-it.ch

## References

1. House, C.M., Frew, I.J., Huang, H.L., Wiche, G., Traficante, N., Nice, E., Catimel, B., and Bowtell, D.D. (2003). A binding motif for Siah ubiquitin ligase. Proc Natl Acad Sci U S A 100, 3101–3106.

2. Bitoun, E., and Davies, K.E. (2005). The robotic mouse: unravelling the function of AF4 in the cerebellum. Cerebellum 4, 250–260.

3. Meyer, C., Hofmann, J., Burmeister, T., Groger, D., Park, T.S., Emerenciano, M., Pombo de Oliveira, M., Renneville, A., Villarese, P., Macintyre, E., et al. (2013). The MLL recombinome of acute leukemias in 2013. Leukemia 27, 2165–2176.

4. Nilson, I., Reichel, M., Ennas, M.G., Greim, R., Knorr, C., Siegler, G., Greil, J., Fey, G.H., and Marschalek, R. (1997). Exon/intron structure of the human AF-4 gene, a member of the AF-4/LAF-4/FMR-2 gene family coding for a nuclear protein with structural alterations in acute leukaemia. Br J Haematol 98, 157–169.

5. Oliver, P.L., Bitoun, E., Clark, J., Jones, E.L., and Davies, K.E. (2004). Mediation of Af4 protein function in the cerebellum by Siah proteins. Proc Natl Acad Sci U S A 101, 14901–14906.

6. Luo, Z., Lin, C., and Shilatifard, A. (2012). The super elongation complex (SEC) family in transcriptional control. Nat Rev Mol Cell Biol 13, 543–547.

7. Jonkers, I., and Lis, J.T. (2015). Getting up to speed with transcription elongation by RNA polymerase II. Nat Rev Mol Cell Biol 16, 167–177.

8. Tang, A.H., Neufeld, T.P., Rubin, G.M., and Muller, H.A. (2001). Transcriptional regulation of cytoskeletal functions and segmentation by a novel maternal pair-rule gene, lilliputian. Development 128, 801–813.

9. Wittwer, F., van der Straten, A., Keleman, K., Dickson, B.J., and Hafen, E. (2001). Lilliputian: an AF4/FMR2-related protein that controls cell identity and cell growth. Development 128, 791–800.

10. Gecz, J., Gedeon, A.K., Sutherland, G.R., and Mulley, J.C. (1996). Identification of the gene FMR2, associated with FRAXE mental retardation. Nat Genet 13, 105–108.

11. Metsu, S., Rooms, L., Rainger, J., Taylor, M.S., Bengani, H., Wilson, D.I., Chilamakuri, C.S., Morrison, H., Vandeweyer, G., Reyniers, E., et al. (2014). FRA2A is a CGG repeat expansion associated with silencing of AFF3. PLoS Genet 10, e1004242.

12. Luo, Z., Lin, C., Woodfin, A.R., Bartom, E.T., Gao, X., Smith, E.R., and Shilatifard, A. (2016). Regulation of the imprinted Dlk1-Dio3 locus by allele-specific enhancer activity. Genes Dev 30, 92–101.

13. Wang, Y., Shen, Y., Dai, Q., Yang, Q., Zhang, Y., Wang, X., Xie, W., Luo, Z., and Lin, C. (2017). A permissive chromatin state regulated by ZFP281-AFF3 in controlling the imprinted Meg3 polycistron. Nucleic Acids Res 45, 1177–1185.

14. Zhang, Y., Wang, C., Liu, X., Yang, Q., Ji, H., Yang, M., Xu, M., Zhou, Y., Xie, W., Luo, Z., et al. (2018). AFF3-DNA methylation interplay in maintaining the mono-allelic expression pattern of XIST in terminally differentiated cells. J Mol Cell Biol.

15. Steichen-Gersdorf, E., Gassner, I., Superti-Furga, A., Ullmann, R., Stricker, S., Klopocki, E., and Mundlos, S. (2008). Triangular tibia with fibular aplasia associated with a microdeletion on 2q11.2 encompassing LAF4. Clin Genet 74, 560–565.

16. Izumi, K., Nakato, R., Zhang, Z., Edmondson, A.C., Noon, S., Dulik, M.C., Rajagopalan, R., Venditti, C.P., Gripp, K., Samanich, J., et al. (2015). Germline gain-of-function mutations in AFF4 cause a developmental syndrome functionally linking the super elongation complex and cohesin. Nat Genet 47, 338–344.

17. Raible, S.E., Mehta, D., Bettale, C., Fiordaliso, S., Kaur, M., Medne, L., Rio, M., Haan, E., White, S.M., Cusmano-Ozog, K., et al. (2019). Clinical and molecular spectrum of CHOPS syndrome. Am J Med Genet A.

18. Urano, A., Endoh, M., Wada, T., Morikawa, Y., Itoh, M., Kataoka, Y., Taki, T., Akazawa, H., Nakajima, H., Komuro, I., et al. (2005). Infertility with defective spermiogenesis in mice lacking AF5q31, the target of chromosomal translocation in human infant leukemia. Mol Cell Biol 25, 6834–6845.

19. Deciphering Developmental Disorders, S. (2015). Large-scale discovery of novel genetic causes of developmental disorders. Nature 519, 223–228.

20. Retterer, K., Juusola, J., Cho, M.T., Vitazka, P., Millan, F., Gibellini, F., Vertino-Bell, A., Smaoui, N., Neidich, J., Monaghan, K.G., et al. (2016). Clinical application of whole-exome sequencing across clinical indications. Genet Med 18, 696–704.

21. Alfaiz, A.A., Micale, L., Mandriani, B., Augello, B., Pellico, M.T., Chrast, J., Xenarios, I., Zelante, L., Merla, G., and Reymond, A. (2014). TBC1D7 mutations are associated with intellectual disability, macrocrania, patellar dislocation, and celiac disease. Hum Mutat 35, 447–451.

22. Delafontaine J., M.A., Liechti R., Kuznetsov D., Xenarios I., Pradervand S. (2016). Varapp: A reactive web-application for variants filtering. bioRxiv preprint.

23. Holla, O.L., Bock, G., Busk, O.L., and Isfoss, B.L. (2014). Familial visceral myopathy diagnosed by exome sequencing of a patient with chronic intestinal pseudo-obstruction. Endoscopy 46, 533–537.

24. Bowling, K.M., Thompson, M.L., Amaral, M.D., Finnila, C.R., Hiatt, S.M., Engel, K.L., Cochran, J.N., Brothers, K.B., East, K.M., Gray, D.E., et al. (2017). Genomic diagnosis for children with intellectual disability and/or developmental delay. Genome Med 9, 43.

25. Takenouchi, T., Yamaguchi, Y., Tanikawa, A., Kosaki, R., Okano, H., and Kosaki, K. (2015). Novel overgrowth syndrome phenotype due to recurrent de novo PDGFRB mutation. J Pediatr 166, 483–486.

26. Richards, S., Aziz, N., Bale, S., Bick, D., Das, S., Gastier-Foster, J., Grody, W.W., Hegde, M., Lyon, E., Spector, E., et al. (2015). Standards and guidelines for the interpretation of sequence variants: a joint consensus recommendation of the American College of Medical Genetics and Genomics and the Association for Molecular Pathology. Genet Med 17, 405–424.

27. Sadedin, S.P., Dashnow, H., James, P.A., Bahlo, M., Bauer, D.C., Lonie, A., Lunke, S., Macciocca, I., Ross, J.P., Siemering, K.R., et al. (2015). Cpipe: a shared variant detection pipeline designed for diagnostic settings. Genome Med 7, 68.

28. Sievers, F., Wilm, A., Dineen, D., Gibson, T.J., Karplus, K., Li, W., Lopez, R., McWilliam, H., Remmert, M., Soding, J., et al. (2011). Fast, scalable generation of high-quality protein multiple sequence alignments using Clustal Omega. Mol Syst Biol 7, 539.

29. Waterhouse, A.M., Martin, D.M.A., Barton, G.J., Procter, J.B., and Clamp, M. (2009). Jalview Version 2—a multiple sequence alignment editor and analysis workbench. Bioinformatics 25, 1189–1191.

30. Santelli, E., Leone, M., Li, C., Fukushima, T., Preece, N.E., Olson, A.J., Ely, K.R., Reed, J.C., Pellecchia, M., Liddington, R.C., et al. (2005). Structural analysis of Siah1-Siah-interacting protein interactions and insights into the assembly of an E3 ligase multiprotein complex. J Biol Chem 280, 34278–34287.

31. Johansson, M.U., Zoete, V., Michielin, O., and Guex, N. (2012). Defining and searching for structural motifs using DeepView/Swiss-PdbViewer. BMC Bioinformatics 13, 173.

32. House, C.M., Hancock, N.C., Moller, A., Cromer, B.A., Fedorov, V., Bowtell, D.D., Parker, M.W., and Polekhina, G. (2006). Elucidation of the substrate binding site of Siah ubiquitin ligase. Structure 14, 695–701.

33. Skarnes, W.C., Rosen, B., West, A.P., Koutsourakis, M., Bushell, W., Iyer, V., Mujica, A. O., Thomas, M., Harrow, J., Cox, T., et al. (2011). A conditional knockout resource for the genome-wide study of mouse gene function. Nature 474, 337–342.

34. Mikhaleva, A., Kannan, M., Wagner, C., and Yalcin, B. (2016). Histomorphological Phenotyping of the Adult Mouse Brain. Curr Protoc Mouse Biol 6, 307–332.

35. Kraft, K., Geuer, S., Will, A.J., Chan, W.L., Paliou, C., Borschiwer, M., Harabula, I., Wittler, L., Franke, M., Ibrahim, D.M., et al. (2015). Deletions, Inversions, Duplications: Engineering of Structural Variants using CRISPR/Cas in Mice. Cell Rep 10, 833–839.

36. Mundlos, S. (2000). Skeletal morphogenesis. Methods Mol Biol 136, 61–70.

37. Sobreira, N., Schiettecatte, F., Valle, D., and Hamosh, A. (2015). GeneMatcher: a matching tool for connecting investigators with an interest in the same gene. Hum Mutat 36, 928–930.

38. Karczewski, K.J., Francioli, L.C., Tiao, G., Cummings, B.B., Alföldi, J., Wang, Q., Collins, R.L., Laricchia, K.M., Ganna, A., Birnbaum, D.P., et al. (2019). Variation across 141,456 human exomes and genomes reveals the spectrum of loss-of-function intolerance across human protein-coding genes. bioRxiv, 531210.

39. Kumar, P., Henikoff, S., and Ng, P.C. (2009). Predicting the effects of coding non-synonymous variants on protein function using the SIFT algorithm. Nat Protoc 4, 1073–1081.

40. Choi, Y., and Chan, A.P. (2015). PROVEAN web server: a tool to predict the functional effect of amino acid substitutions and indels. Bioinformatics 31, 2745–2747.

41. Adzhubei, I.A., Schmidt, S., Peshkin, L., Ramensky, V.E., Gerasimova, A., Bork, P., Kondrashov, A.S., and Sunyaev, S.R. (2010). A method and server for predicting damaging missense mutations. Nat Methods 7, 248–249.

42. Schwarz, J.M., Cooper, D.N., Schuelke, M., and Seelow, D. (2014). MutationTaster2: mutation prediction for the deep-sequencing age. Nature Methods 11, 361.

43. Pérez-Palma, E., May, P., Iqbal, S., Niestroj, L.-M., Du, J., Heyne, H., Castrillon, J., O’Donnell-Luria, A., Nürnberg, P., Palotie, A., et al. (2019). Identification of pathogenic variant enriched regions across genes and gene families. bioRxiv, 641043.

44. Brown, S.D., and Moore, M.W. (2012). The International Mouse Phenotyping Consortium: past and future perspectives on mouse phenotyping. Mamm Genome 23, 632–640.

45. Fietz, S.A., Lachmann, R., Brandl, H., Kircher, M., Samusik, N., Schroder, R., Lakshmanaperumal, N., Henry, I., Vogt, J., Riehn, A., et al. (2012). Transcriptomes of germinal zones of human and mouse fetal neocortex suggest a role of extracellular matrix in progenitor self-renewal. Proc Natl Acad Sci U S A 109, 11836–11841.

46. Moore, J.M., Oliver, P.L., Finelli, M.J., Lee, S., Lickiss, T., Molnar, Z., and Davies, K.E. (2014). Laf4/Aff3, a gene involved in intellectual disability, is required for cellular migration in the mouse cerebral cortex. PLoS One 9, e105933.

47. Luo, Z., Lin, C., Guest, E., Garrett, A.S., Mohaghegh, N., Swanson, S., Marshall, S., Florens, L., Washburn, M.P., and Shilatifard, A. (2012). The super elongation complex family of RNA polymerase II elongation factors: gene target specificity and transcriptional output. Mol Cell Biol 32, 2608–2617.

48. Nakamura, M., Matsuda, Y., Higo, M., and Nishimura, G. (2007). A family with an autosomal dominant mesomelic dysplasia resembling mesomelic dysplasia Savarirayan and Nievergelt types. Am J Med Genet A 143A, 2079–2081.

49. Bonafe, L., Cormier-Daire, V., Hall, C., Lachman, R., Mortier, G., Mundlos, S., Nishimura, G., Sangiorgi, L., Savarirayan, R., Sillence, D., et al. (2015). Nosology and classification of genetic skeletal disorders: 2015 revision. Am J Med Genet A 167A, 2869–2892.

50. Tuysuz, B., Zeybek, C., Zorer, G., Sipahi, O., and Ungur, S. (2002). Patient with the mesomelic dysplasia, Nievergelt syndrome, and cerebellovermian agenesis and cataracts. Am J Med Genet 109, 206–210.

51. Savarirayan, R., Cormier-Daire, V., Curry, C.J., Nashelsky, M.B., Rappaport, V., Rimoin, D. L., and Lachman, R.S. (2000). New mesomelic dysplasia with absent fibulae and triangular tibiae. Am J Med Genet 94, 59–63.

52. Lefevre, L., Omeiri, H., Drougat, L., Hantel, C., Giraud, M., Val, P., Rodriguez, S., Perlemoine, K., Blugeon, C., Beuschlein, F., et al. (2015). Combined transcriptome studies identify AFF3 as a mediator of the oncogenic effects of beta-catenin in adrenocortical carcinoma. Oncogenesis 4, e161.

53. Zhong, Z., Ethen, N.J., and Williams, B.O. (2014). WNT signaling in bone development and homeostasis. Wiley Interdiscip Rev Dev Biol 3, 489–500.

54. Tufan, F., Cefle, K., Turkmen, S., Turkmen, A., Zorba, U., Dursun, M., Ozturk, S., Palanduz, S., Ecder, T., Mundlos, S., et al. (2005). Clinical and molecular characterization of two adults with autosomal recessive Robinow syndrome. Am J Med Genet A 136, 185–189.

55. Kimura, R., Watanabe, C., Kawaguchi, A., Kim, Y.I., Park, S.B., Maki, K., Ishida, H., and Yamaguchi, T. (2015). Common polymorphisms in WNT10A affect tooth morphology as well as hair shape. Hum Mol Genet 24, 2673–2680.

56. Melko, M., Douguet, D., Bensaid, M., Zongaro, S., Verheggen, C., Gecz, J., and Bardoni, B. (2011). Functional characterization of the AFF (AF4/FMR2) family of RNA-binding proteins: insights into the molecular pathology of FRAXE intellectual disability. Hum Mol Genet 20, 1873–1885.

