## Supplementary material for "Variants in the degron of *AFF3* cause a multi-system disorder with mesomelic dysplasia, horseshoe kidney and developmental and epileptic encephalopathy": Table S1

**Table S1. Phenotype of individuals with *AFF3* variants**

| Individual | Proband 1 | Proband 2 | Proband 3 | Proband 4 | Proband 5 | Proband 6 | Proband 7 | Proband 8 | Proband 9 | Proband 10 | Deletion proband* |
|---|---|---|---|---|---|---|---|---|---|---|---|
| **Age [years]** | 17 | 3 years | 18 | 21 | 5 | 9 | 8 | 8 | 8 | 11 | 4 months |
| **Sex** | M | F | M | F | F | F | F | F | M | M | F |
| **AFF3 variant** | p.(A258S) | p.(A258S) | p.(A258T) | p.(A258T) | p.(A258T) | p.(A258T) | p.(A258T) | p.(A258T) | p.(A258V) | p.(V260G) | 500kb deletion |
| **Inheritance** | *De novo* | *De novo* | *De novo* | *De novo* | *De novo* | *De novo* | *De novo* | *De novo* | *De novo* | *De novo* | *De novo* |
| **Neurodevelopmental anomalies** | | | | | | | | | | | |
| **Severe DD/ID** | + | + | + | + | + | + | + | + | + | + | + |
| **Epilepsy** | Generalized tonic-clonic seizures (onset at 5 years), nocturnal, treatment with Keppra and Micropakine | Multifocal epileptiform discharges in bilateral posterior quadrant, no clinical seizures. | Generalized, tonic-clonic seizures (onset at 5 months), initially controlled with phenobarbital, developed drug-resistance, partially controlled with carbamazepine, clobazam, nitrazepam and phenobarbital | Generalized, tonic-clonic seizures | Generalized, tonic-clonic seizures | Complex partial with secondary generalization | Generalized, tonic-clonic seizures | Generalized (onset at 3 years), tonic-clonic seizures, inefficient treatments but remission since 6 years 9 months | - | Generalized, tonic-clonic seizures | Myoclonic jerks, convulsions |
| **Muscle tone** | Limb hypertonia, spastic tetraparesis | Generalized hypotonia | Axial hypotonia, peripheric hypertonia | Limb hypertonia | Hypotonia | Axial hypotonia, peripheric hypertonia | Hypotonia | Hypotonia | Normal | Hypotonia | N/A |
| **Vision and hearing** | Normal | Cortical visual impairment, hyperopic refractive error and small angle intermittent strabismus | Normal | Myopia, strabismus | Strabismus | Central vision loss due to occipital impairment, central progressive hearing loss | Normal | Strabismus inconstant | Conductive hearing loss (resolved), vision not yet formally assessed | Strabismus | N/A |
| **Brain MRI** | Enlargement of the ventricular system and peri-cerebral spaces, thin and irregular appearance of the corpus callosum | Partial agenesis of the corpus callosum, forshortened and undersulcated frontal lobes, small cerebellar vermis with mega cisterna magna and wide sylvian fissures | Pachygyria of frontal lobes, abnormal opercularization of insulae with thicker insular cortices, wide Sylvian fissures, cerebral atrophy, brainstem hypoplasia, increased volume of the trigones and occipital horns of the lateral ventricles | N/A | Cerebral atrophy with possible brainstem hypoplasia | Prominence of CSF spaces | Cerebral atrophy | Cerebral atrophy, pachygyria of frontal lobes | Normal | N/A | Brain atrophy, ventriculomegaly |
| **Other** | Stereotypic movements | - | - | - | - | - | - | - | Autism and ADHD | - | - |

| | | | | | | | | | | | |
|---|---|---|---|---|---|---|---|---|---|---|---|
| **Craniofacial features** | | | | | | | | | | | |
| **Microcephaly** | + (plagiocephaly) | - | + | - | + | - (brachycephaly) | + | + | + | + | -(dolicocephaly) |
| **Nose** | Small nose, anteverted nares | Prominent columella | Large nose with bulbous nasal tip and low hanging columella | Large nose with bulbous nasal tip and low hanging columella | Small nasal tip | Bulbous nasal tip, low hanging columella with low insertion | Large nose with bulbous nasal tip and low hanging columella | Large nose with bulbous nasal tip and low hanging columella | Normal | N/A | N/A |
| **Short philtrum** | - | - | + | + | - | + (smooth) | + | + | Smooth philtrum | N/A | N/A |
| **Mouth** | Wide mouth with square upper lip | Wide mouth, downturned corners, thin upper lip | Wide mouth with square upper lip | Wide mouth with square upper lip | Wide mouth with square upper lip | Wide mouth with downturned corners, thick lower lip vermillion | Prominent upper lip | Wide mouth | Thin upper vermillion, mild ankyloglossia (snipped), high arched palate | Ankyloglossia | N/A |
| **Teeth and gum abnormalities** | Small, gingival hypertrophy | Small, widely spaced, bruxism | + | + | Small, widely spaced | + | - | Widely spaced | Hypomineralisation | + | N/A |
| **Chin** | Prognathism | Normal | Micrognathia | Triangular chin | Normal | Micrognathia | Mild micrognathia | Prognathism | Micrognathia | N/A | N/A |
| **Synophrys and hypertrichosis** | + | - | + | + | + | + | + | + | + | N/A | N/A |
| **Other** | Hypertelorism, short neck | Full cheeks, mild facial asymmetry | - | - | - | Long palpebral fissures, low-set and posteriorly rotated large ears with a simple helix, facial asymmetry | Long palpebral fissures | Long palpebral fissures, low set ears, mild facial asymmetry, gingival hyperplasia diabetes | Slight metopic prominence | - | Short palpebral fissures, low set ears, short neck |
| **Skeletal abnormalities** | | | | | | | | | | | |
| **Mesomelic dysplasia** | - | Lower limbs | 4 limbs | 4 limbs | 4 limbs | 4 limbs | 4 limbs | 4 limbs | Mild lower limbs | Lower limbs | 4 limbs |
| **Arms and legs** | Bilateral elbow dislocation | Bilateral fibular agenesis, short and curved tibia, bilateral Syme amputations with resection of cartilaginous fibular anlage and bilateral tibial osteotomies for angular deformity correction, fitted with bilateral lower extremity prosthetics at 2 years 3 months. | Madelung deformity, slender limb bones, fibular hypoplasia/agenesis | Short ulna and radius, radial head dislocation/subluxation, styloid process of ulna on radius, carpal coalition, hypoplastic femora, short and curved tibia with metaphyseal flaring, mid tibial dimples, deviated knees, hypoplastic and gracile fibula | Short and thick ulna, slightly shortened radius with convex distal end bilaterally, dislocation of right radial head, short and curved tibia, extremely short rectangular fibula | Short fibula, discoid meniscus, limited knee extension | Limited supination, radial head dislocation/subluxation, hypoplastic fibula | Short humerus, hypoplastic short fibula | Mild lateral bowing of both radii, normal bone age | Bowed radii, unilateral bowed ulna, shortened ulna, abnormal radial diaphysis, bowed and angulated tibias, hypoplastic fibula | Radial head dislocation/subluxation, slightly short radius and ulna, short and dysplastic triangular tibias, fibular agenesis |
| **Hands and feet** | Limited pronosupination, bilateral camptodactyly, edema of back-hands and -feet, bilateral simian creases, tapered fingers, dorsum pedis edema, small toes, hypoplasia of | Right single transverse and left bridged palmar crease, bilateral hypoplastic 4$^{th}$ metatarsals, absence of the 5$^{th}$ ray and phalanges of lateral toes, 4 splayed toes | Hypoplastic talipes, fusion of tarsal bones | Carpal coalition, small feet, hypoplastic left 5$^{th}$, metatarsal synostosis | Talus ossified in hindfoot, one ossified bone in midfoot (cuneiform), missing one lateral ray in left foot, 4$^{th}$-5$^{th}$ right metatarsal synostosis | Soft tissue syndactyly of fingers 3$^{rd}$-4$^{th}$, small feet, pes planus, 2$^{nd}$ toe overlapping hallux bilaterally | Limited supination, pes planus, broad toe tips | Transverse palmar crease, limited pronosupination, proximal deviation of thumbs, small feet, absent calcanei, broad 1$^{th}$ toes, polydactyly, cutaneous | Very broad feet. Bilateral hind foot varus deformity, mild metatarsus adductus. misshaped and large talus, cuboid and calcaneum, mild shortening and Y | Wide distal radial metaphyses, oligodactyly: 2 tarsal bones on each foot, absent/hypoplastic calcanei, 3 metatarsals, 3 associated phalanges, 1 phalanx not | Small equinovalgus feet, oligodactyly: 4 toes on 1 foot, 5 on the other, abnormally spaced |

| | | | | | | | | | | | |
|---|---|---|---|---|---|---|---|---|---|---|---|
| | distal phalanges, ungual hypoplasia, neonatal arthrogryposis | | | | | | | process on the side of the 5$^{th}$ finger and cutaneous syndactyly 3$^{th}$-6$^{th}$ toes on left foot, four metatarsals and partial syndactyly 3$^{th}$-4$^{th}$ toes on right foot | shaped fusion of the left 4$^{th}$ and 5$^{th}$ metatarsals, hyperplastic/dysplastic toenails, hand Xrays normal | associated with a metatarsal bone | |
| **Ribs and spine** | Scoliosis | 13 rib-bearing thoracic-type vertebrae and 5 lumbar type vertebrae, hypoplastic L1 with focal kyphosis | Severe scoliosis, C2-C3 vertebral fusion, L5-S1 vertebral cleft | Scoliosis | Bilateral cervical ribs | Scoliosis, incomplete coronal cleft of T9 and T12 vertebrae, low lying spinal cord, termination of conus medullaris at upper border of L3 | Pectus excavatum | Scoliosis, fusion 1$^{th}$-2$^{th}$ ribs, sacral sinus | 6 lumbar vertebrae, 13 pairs of ribs, sacral dimple | Scoliosis, cervical ribs, anterior superior vertebral notching, tethered cord | Sacral sinus |
| **Hips and pelvis** | Bilateral coxa valga, dislocation of the hips | Normal | Normal | Bilateral coxa valga with hypoplastic ilia, hip dislocation | Coxa valga | Bilateral coxa valga | Hip dislocation | Coxa valga, hip dysplasia | Normal | Coxa valga, unilateral hip dysplasia | N/A |
| **Osteopenia** | N/A | + | + (osteoporosis) | + | - | + | + (osteoporosis) | - | - | + | N/A |
| **Additional features** | | | | | | | | | | | |
| **Horseshoe kidney** | - | + | - | + | + | + | - | + | + | + | + |
| **Gastro-intestinal symptoms** | N/A | GERD, dysphagia, gastrostomy tube dependent, concerns for esophageal dysmotility ± abnormal gastric accommodation, abnormal gastric emptying with no evidence of small intestinal dysmotility | GERD, constipation | GERD, constipation | Constipation, swallowing difficulties, percutaneous endoscopic gastrostomy | GERD, gastrojejunostomy tube dependent, chronic constipation, hiatal hernia, pancreatitis | Constipation | Constipation, anal dystopia | Constipation, feeding difficulties due to floppy larynx and adenotonsillar hypertrophy, frequent obstruction | GERD, constipation | Colon malrotation |
| **Failure to thrive** | + | - | + | + | + | + | + | + | + | + | + |
| **Respiratory problems** | Apnea | Multicompartmental respiratory disease (upper airway obstruction, lower airway obstruction, ineffective mucociliary clearance, restrictive lung disease, aspiration and pneumonia), moderate to | Neonatal respiratory distress, recurrent pneumonia, frequent hiccups improved with carbamazepine, respiratory arrest leading to death at 21 years | - | Nightly desaturations treated with CPAP from 3 years of age | - | - | - | Severe obstructive sleep apnea, adenotonsillectomy at 3 years | - | Recurrent apnea, respiratory arrest leading to death at 4 months |

| | | | | | | | | | |
|---|---|---|---|---|---|---|---|---|---|
| | | severe mixed sleep apnea, severe laryngomalacia status post supraglottoplasty at 18 months, cough assist and inhaled steroid and bronchodilator and supplemental oxygen with sleep, tonsillectomy and adenoidectomy planned | | | | | | | |
| **Other** | Bilateral cryptorchidism, bicuspid aortic valve | History of bilateral vesicoureteral reflux, grade II | | Menstrual cycle perturbations | | | | Popliteal pterygium | Short stature |

**Footnote:** AFF3 variants are described according to RefSeq NP_002276.2. ***Steichen-Gersdorf *et al* (2008)**[15]
**Abbreviations:** DD= developmental delay, ID= intellectual disability, N/A = not available, ADHD= Attention deficit hyperactivity disorder, GERD= gastroesophageal reflux disease
