## Supplementary material for "Variants in the degron of *AFF3* cause a multi-system disorder with mesomelic dysplasia, horseshoe kidney and developmental and epileptic encephalopathy": Figure S1

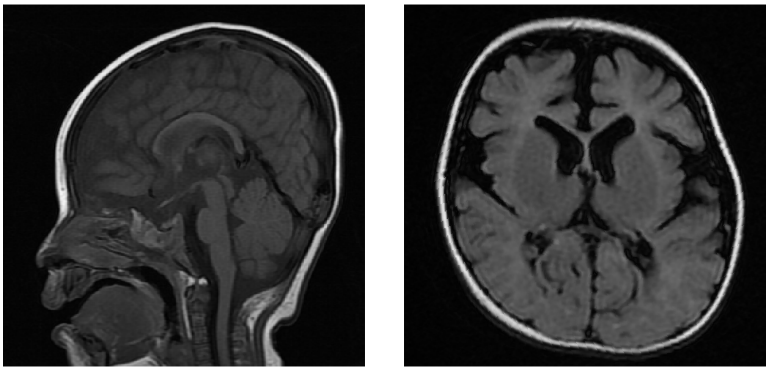

**Figure S1:** Brain MRI of proband 7 carrying a *de novo* variant in *AFF3*
FLAIR (right) and T1 (left) at 9 months old
